# Evidence against the microbicidal action of neutrophil extracellular traps (NETs)

**DOI:** 10.1101/389593

**Authors:** F. Semplici, A. W. Segal

## Abstract

Neutrophil extracellular traps (NETs) are fibrillary structures composed of extruded nuclear chromatin decorated with granule proteins (mostly neutrophil elastase, cathepsin G and myeloperoxidase). It has been reported that NETs are able to kill bacteria and fungi based upon the observation that smaller number of organisms are obtained in plating assays after they are incubated with NETs than if the DNA is pre-digested with DNase. It is possible that the microbial killing is apparent rather than real, and occurs because the organisms are aggregated on the DNA structure, and that the plating assay results were simply misinterpreted. The present study shows that digestion of DNA after incubation of NETs with the microbes restores their numbers to preincubation levels indicating that the apparent killing is an artefact of the assay.

## Introduction

Neutrophil extracellular traps (NETs) have been proposed as a mechanism for killing fungi and bacteria (1–3). NETs are extrusions of DNA from neutrophils, which is released from cells stimulated with phorbol myristate acetate (PMA) and various other agents, including lipopolysaccharides (LPS) and interleukin 8 (IL8). The mechanism by which they are produced appears to be through translocation of neutrophil elastase from cytoplasmic granules to the nucleus (4), digestion of histones and other nuclear proteins, decondensation of chromatin, and the release of DNA from the nucleus (3), and also from the mitochondria under certain circumstances (5).

Microbes have been shown to attach to the DNA mesh (1), as do the neutrophil granule enzymes, (1) and it has been proposed that they are killed by the enzymes in this location (1). The assay used to demonstrate this effect depends upon the production of NETs in plastic wells, the incubation of these NETs with microbes after which the organisms are harvested from the wells and their numbers determined by colony counts. The specific effect of the NETs is determined by predigesting the DNA in the control wells. A potential problem that could arise from the mixture of DNA and microbes is that the attachment of the organisms to the DNA could impede their dispersal, resulting in a reduced number of colonies, giving a false impression of microbial killing.

One way of countering the aggregation of microbes on DNA would be a brief treatment with DNase after the incubation but before the plating stage. This experiment has already been performed and demonstrated that the apparent killing of the organisms could be completely reversed by DNase (6, 7). If the conclusions of this study are correct, this is important because of the attention NETs have attracted, based upon the belief that they kill microbes. In addition, concerns have been raised as to the microbicidal action of these structures (8, 9). The present study has been undertaken to determine whether or not the addition of DNase to the suspension of cells and microbes after the incubation period reverses the apparent microbicidal effect of the NETs.

## Materials and Methods

### Reagents

Histopaque 1119 (Cat # 11191-100ML), Albumin from Human Serum (Cat # A1653-5G), Human Serum (Cat # H4522), Deoxyribonuclease I (Cat # D4527-10KU), Hepes (Cat # H0887-100ML) and Phorbol 12-Myristate 13-Acetate (PMA) (Cat # P1585-1MG) were from Sigma; Percoll (Cat # 17-0891-02) was from GE Healthcare, Yeast Extract (Cat # LP0021B) was from Oxoid, Peptone (Cat # 211677) was from Becton Dickinson, Sytox Green (Cat # S7020) and RPMI 1640 medium without phenol red (Cat # 11835063) were from ThermoFisher Scientific. Proteinase K (Cat #3115801001) was from Roche.

### Bacterial Strains and Growth Conditions

*C. albicans* (Cat ATCC®10231) vitroids were from Sigma, *E. coli* strain DH5 alpha was from New England Biolabs (Cat # C2987), and *S. tiphimurium* expressing mCherry cDNA was obtained from F. Buss (CIMR, University of Cambridge, UK).

*C. albicans*, E. coli and S.tiphimurium were grown overnight, respectively in YPD medium at 30°C, and in LB broth at 37°C, then washed in PBS before being added to the neutrophils.

### Human Neutrophil Isolation

Neutrophils were isolated from whole peripheral blood obtained with informed consent from healthy donors as described (10). Briefly, 20 ml of blood were separated first on Histopaque 1119 and then on Percoll discontinuous gradient, followed by a haemolytic shock.

### NET formation and Bacterial Infection

Neutrophils (1×10^6^/well) were plated in a 24-well plate in RPMI without phenol red, supplemented with 10 mM Hepes, and incubated at 37°C, 5% CO_2_ for 1 hour, prior to stimulation with 100 nM PMA for 4 hours, to allow NET production. At the end of this time, freshly grown microorganisms (mid-log phase) were added to the neutrophils in a ratio of 1:1, in the presence or absence of DNase I (100 U/ml), and the cells were further incubated as above for 1 hour. Where indicated, at the end of the 1 hour incubation, DNase I (100U/ml) either active or heat inactivated (HI DNase) (at 75°C for 20 minutes), was added to the wells where it had not been previously added, for a further 10 minutes. All wells were then treated with Proteinase K (at a final concentration of 20 μg/ml) to release the bacteria from the plastic surface. Finally, the plate was kept on ice while the contents of the wells were collected after scraping the well with a cell scraper. Serial dilutions of the contents of each well were plated on Yeast extract Peptone Dextrose (YEPD) (for *C. albicans*) or Lysogeny Broth (LB) (for *E. coli* and *S. typhimurium*) plates and colonies counted. Each experiment was performed three times in which each incubation was performed in duplicate.

### Statistical analysis

Results are the means ± standard error. Data were analysed by one-way ANOVA, followed by Bonferroni. P< 0.05 was considered significant.

### Ethics

Ethical and research governance approval was provided by The National Research Ethics Service London Surrey Borders Committee (10/H0906/115) and the University College London Research Ethics Committee (6054/001). Informed consent was obtained from all participants.

## Results and Discussion

Human neutrophils stimulated with 100 nM PMA for 4 hours resulted in the production of abundant NETs, as previously described (10).

Figure 1 A shows that in the absence of DNase the NETs reduced the numbers of *C. albicans* by more than 75%. If, however, the preparations were briefly incubated with DNase after the incubation period during which killing was considered to have taken place, which we confirmed abolished the NETs, the numbers of *Candida* were restored. This effect was not seen with heat inactivated DNase. These results indicate that killing had not occurred and that the impression that it had was due to sequestration on the DNA mesh.

**Figure 1.**
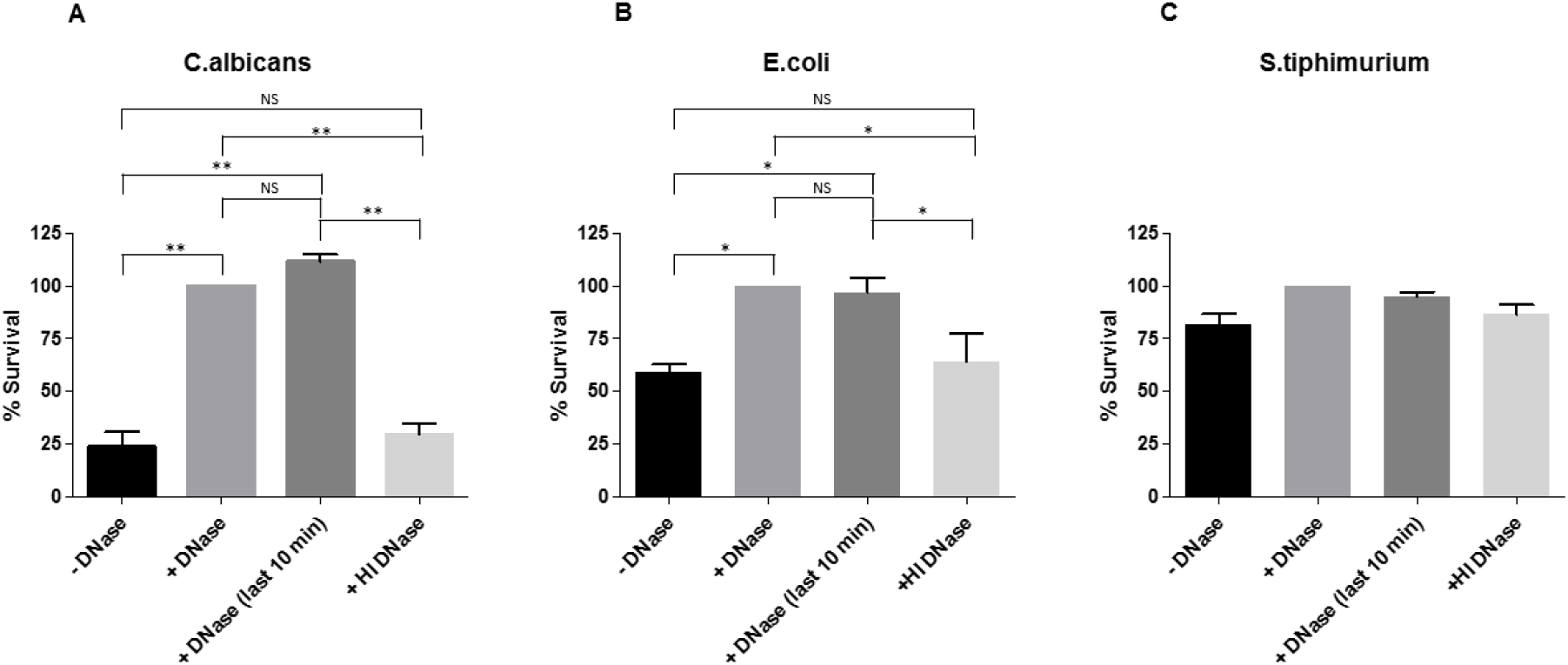
Survival of *Candida albicans*, *Escherichia coli* and *Salmonella typhimurium* after incubation with NETs, in the absence or presence of DNase. Survival of (A) *C. albicans*, (B) *E. coli* and (C) *S. typhimurium* after incubation with NETs in the absence and presence of DNase from the outset. The apparent killing of the microbes by the NETs was reversed at the end of the incubation by a brief incubation of wells that had not previously contained DNase, with active [+ DNase (last 10 min] but not heat inactived DNase (HI DNase), indicating that true killing of the microbes by NETs had not taken place. NS, not significant (P>0.05); * P<0.005 and ** P<0.001 (one-way ANOVA followed by Bonferroni’s)

When we used *E. coli* as the test organism the microbicidal activity of the NETs (Figure 1 panel B) we observed that the number of organisms were again reduced by NETs in the absence of DNase but only to 55% of the initial inoculate, and once again the numbers were returned to the original levels by treatment with active, but not inactivated, DNase.

As shown in panel C of Figure 1, there was little apparent “killing” of *S. typhimurium*.

The results of the experiments described here agree with those described previously (6, 7) and clearly demonstrate that the addition of DNase to the preparations containing cells, DNA and microbes reverses the reduction in the numbers of colonies that give rise to the impression of microbial killing by the NETs. DNase could not reverse true microbial death in the ten minutes of the final incubation and the results can only mean that the numbers of microbes were increased by more efficient dispersion as a result of the removal of the DNA to which they were attached, as illustrated in Figure 2.

**Figure 2.**
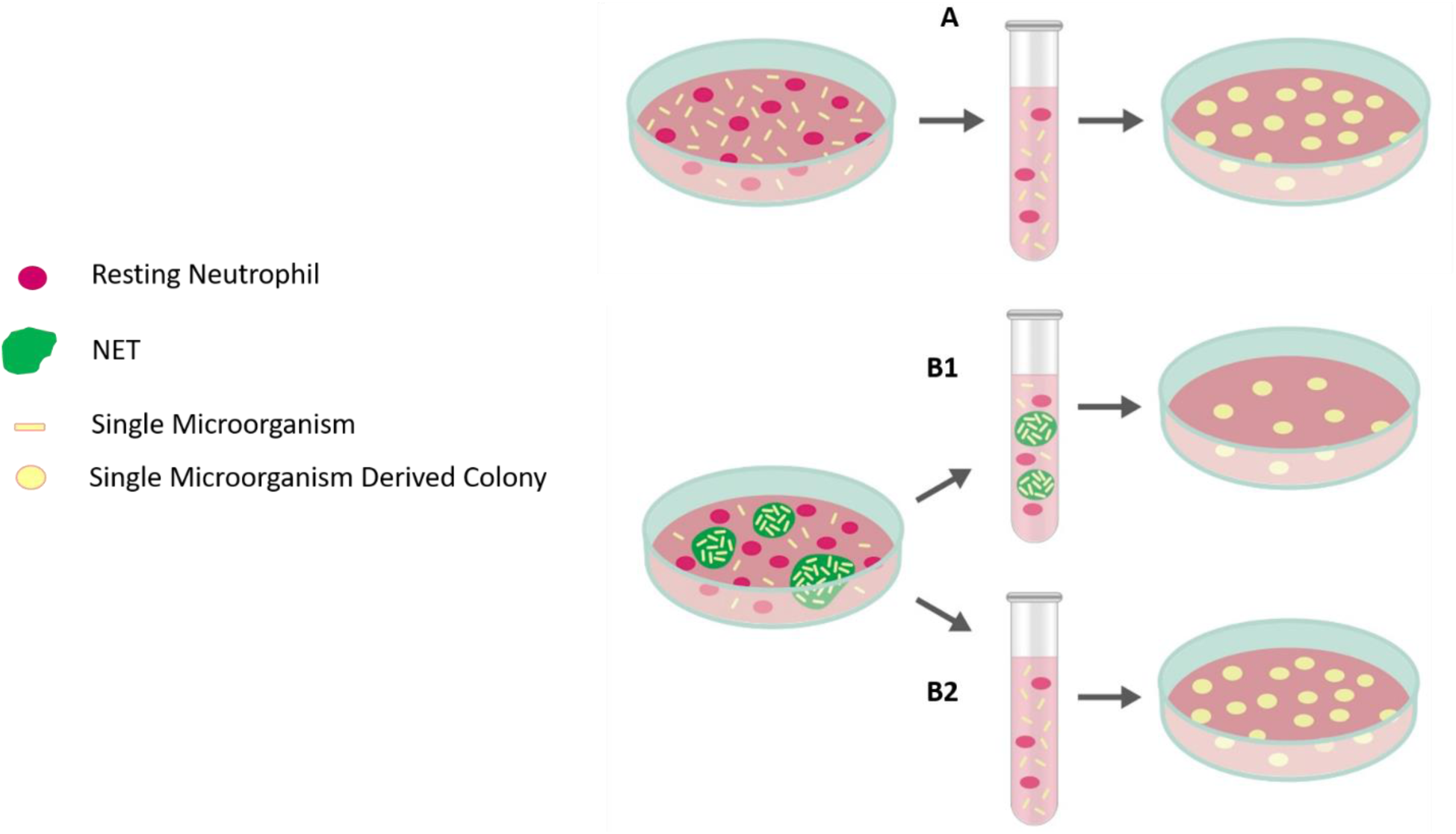
Schematic representation of the proposed mechanism by which NETs appear to kill microbes. In A, DNAse is added to the incubation with NETs before the addition of microbes. In B1 no DNAse is added, whereas in B2 DNAse is added after the incubation of microbes with NETs and before plating. The observed increase in the colony counts between B1 and B2 must indicate that the DNAse is dispersing the microbes by lysing the macrostructure of the DNA that is derived from the NETs.

The variation in the reduction in the numbers of colonies produced by the different organisms when incubated with NETs is probably a reflection of the affinities with which they attach to the DNA filaments.

It has been proposed that neutrophil NETs might play an important role in physically containing infecting microbes (11). They might well add to the network of fibrin and thrombin fibres that are deposited at sites of infection, a process termed “immunothrombosis” (12, 13), as well as many other sources of extracellular DNA (14), that encompass, and in some cases shield and nourish the organisms.

Conclusions arising from those studies (1–3) that demonstrate the apparent killing of microbes by NETs require reevaluation.

### Authorship

F. S. carried out the experimental work and contributed to the preparation of the manuscript.

A.W.S. contributed to the experimental design, interpretation and preparation of the manuscript.

## Acknowledgments

We thank the Medical Research Council for support.

## Abbreviations

NETs: Neutrophil Extracellular Traps
PMA: Phorbol 12-Myristate 13-Acetate
LPS: Lipopolysaccharides
DNase: Deoxyribonuclease
HI: Heat Inactivated
YEPD: Yeast Extract Peptone Dextrose
LB: Lysogeny Broth
CFU: Colony Forming Unit

